# A Game-Theoretic Model for Co-Adaptive Brain-Machine Interfaces

**DOI:** 10.1101/2020.12.11.421800

**Authors:** Maneeshika M. Madduri, Samuel A. Burden, Amy L. Orsborn

## Abstract

Co-adaptation in brain-machine interfaces (BMIs) can improve performance and facilitate user learning. We propose and analyze a mathematical model for co-adaptation in BMIs. We model the brain and the decoder as strategic agents who seek to minimize their individual cost functions, leading to a game-theoretic formulation of interaction. We frame our BMI model as a potential game to identify stationary points (Nash equilibria) of the brain-decoder interactions, which correspond to points at which both the brain and the decoder stop adapting. Assuming the brain and the decoder adapt using gradientbased schemes, we analytically show how convergence to these equilibria depends on agent learning rates. This theoretical framework presents a basis for simulating co-adaption using dynamic game theory and can be extended to tasks with multiple dimensions and to different decoder models. This framework can ultimately be used to inform adaptive decoder design to shape brain learning and optimize BMI performance.

## I. INTRODUCTION

### A. Motivating Background

Uncovering the neural principles of learning in the brain is key to developing high-performing neural interfaces [1], [2]. Closed-loop Brain-Machine Interfaces (BMIs) define an input-output relationship in the brain and can provide insight into fundamental neural mechanisms underlying sensorimotor learning [3]. Motor BMIs use a *decoder* to convert neural data to a control signal, such as moving a computer cursor. Closed-loop BMIs form a closed control loop via visual feedback, creating novel sensorimotor systems that can be manipulated to study neural functions [4].

Learning in BMI can lead to the formation of a *stable cortical map*, similar to natural motor learning [5]. Stable cortical maps are accompanied by stable BMI task performance, such as a consistent cursor trajectory to the target with low variability and other signatures of skillful motor control. Recent research has shown that *closed-loop decoder adaptation* (CLDA)–algorithms to modify decoder parameters during closed-loop BMI operation—are potentially able to guide user learning to facilitate the emergence of stable cortical maps [6], [7]. CLDA algorithms have been designed to improve BMI performance and decrease variability in decoder estimates [8], [9]. Designing CLDA algorithms that can optimize brain learning is an open challenge. As an inherently real-time process, CLDA requires online BMI validation. However, testing a wide range of CLDA algorithms through experimentation is impractical. To constrain the space of testable CLDA design parameters and reduce experimental load, we propose and analyze a mathematical model for closed-loop co-adaptive BMIs.

### B. Related Work

Existing BMI models draw upon control theory to conceptualize the brain and decoder as systems adapting according to specified learning processes and subject to cost functions [10] –[12]. Two of these models [11], [12] identify coadaptation strategies using optimization principles to propose adaptive decoder paradigms. The brain and the decoder in these models are treated as as two distinct stochastic optimal control systems [11] or as two systems optimizing a joint cost function [12]. The learning dynamics of co-adaptation in these models either propose a stochastic coordinate descent, where the decoder learns to anticipate the brain’s adaptation [11] or a stochastic gradient descent with an online leastsquares estimator, where the brain and the decoder estimate each other’s parameters to form internal approximations of the other learner [12]. The proposed optimal brain-decoder solutions for each model are identified through either mathematical analysis [11] or computational simulations [12].

### C. Our Contributions

The purpose of this work is to model the interaction between brain and decoder in a co-adaptive BMI. Our contributions to the existing literature are: (a) formulating the two-learner problem for motor BMI in a game-theoretic framework; (b) deriving *stationary points* (Nash equilibria) of the game; and (c) analyzing convergence of a model for co-adaptation to these stationary points. In particular, we adopt the perspective that the brain and decoder are two agents playing a *dynamic game* [13], [14], and apply techniques from game theory to study the asymptotic dynamics of co-adaptation. We use a *potential game* formulation to derive stationary points (Nash equilibria), which points are our model’s surrogates for *stable cortical maps* observed in experiments. Assuming the brain and decoder adapt using (stochastic) gradient descent, we assess convergence to stationary points. Finally, we corroborate our mathematical analyses using numerical simulations.

## II. METHODS

### A. Mathematical Model for Motor BMI

Fig. 1 displays each component and corresponding symbolic representation. Signals are denoted with lower-case non-bolded symbols (e.g. *x*) and parameterizations of the transformations are **bolded** and capitalized (e.g. **X**). The BMI session is a series of trials, with each trial being represented as a new target presented the user. Time is indexed by the subscript trial number *t* = 1, 2,…, dimension of the BMI task is indexed by the subscript *d* = 1, 2,… and neurons are indexed in a population of size *N* by the superscript *n* ∈ {1,…,*N*). We will suppress the sub- and/or superscripts when they are clear from context.

**Fig. 1.**
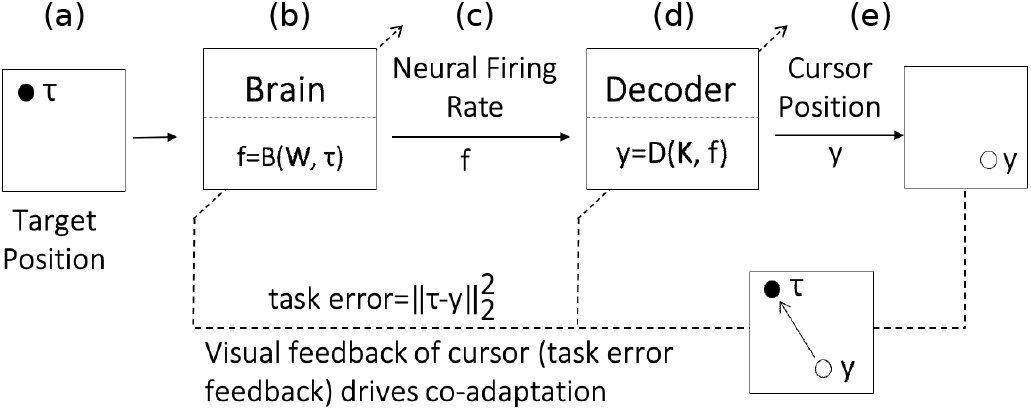
Diagram of the BMI model showing the forward model of the brain and the decoder as well as the visual feedback error of the cursor (task error feedback) that drives the co-adaptation of the brain and the decoder [15]. Based on prior BMI models [10], [11], the elements of this model are: (a) the target presented to the user, (b) the user, (c) the firing rates (recorded as time-binned intracortical neural spiking activity) from a population of neurons, (d) the decoder, (e) the cursor position at the output of the decoder.

#### Target Position

The task presented in this BMI model is a center-out cursor control task–e.g. [6], [7]. The target position is at *τ*.

#### Brain and Firing Rates

The brain is represented as a linear transformation of the target *τ* by a matrix representing the brain’s parameters **W** (1) [10], [12].

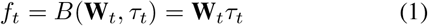

where 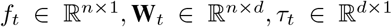. At each new trial *t*, the firing rates are updated and are inputs to the decoder (2).

#### Decoder and Cursor Position

The decoder algorithm in this model is represented as a Wiener filter, a commonly-used experimental decoder that has also been implemented in prior models–e.g. [10], [12]. The Wiener filter was used in this model for simplicity of analysis, but the Kalman Filter, which is a state-based estimator, has been used in co-adaptive BMI experiments–e.g. [6], [7]. The decoder transforms firing rates, *f*, to a *d*-dimensional cursor position, *y*, by multiplying the firing rates by the decoder’s parameters, **K** (2).

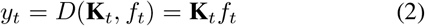

where 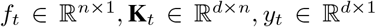.

### B. Brain and Decoder Cost Functions

For each trial, the task error (3) is calculated as the quadratic loss of the cursor and target positions [10], [12]:

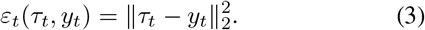

The brain and the decoder parameters, **W** and **K**, update at each trial iteration using a gradient-based scheme detailed in II-C. From a game-theoretic lens, this model can treat the brain and the decoder as two agents in a 2-player game. Each agent, the brain or the decoder, is trying to minimize its own cost function through co-adaptation. The cost function of the brain and decoder are:

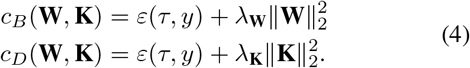

The brain and the decoder cost functions both include regularization terms: 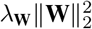 for the brain and 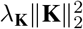 for the decoder. The parameters *λ*_**W**_ and *λ*_**K**_ are scaling factors on the regularization terms and, as we will show later in this paper, play crucial roles in ensuring the convergence of the cost functions. These regularization terms serve to penalize the effort of the brain and the decoder.

### C. Game-Theoretic Stationary Points

#### 1) Defining a Potential Function

With the mathematical model described in II-A, we can reformulate the brain-decoder interactions as a *potential game*. In a potential game, the action of all agents can be expressed with a single function called the potential function. If there exists a continuously differentiable function *ϕ*: 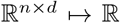 such that 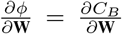 and 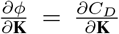, this function *ϕ* can be considered to be a potential function [16]. This highlights an important assumption: the cost functions of the brain and the decoder are differentiable. The brain and the decoder cost functions satisfy the conditions of a potential game and can be thus characterized with the following potential function:

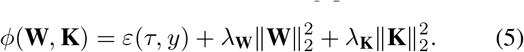

This potential function matches the two-learner system cost function proposed by a prior model [12]. The potential function ϕ tracks the changes of both players, the brain and the decoder in this case, and reformulates co-adaptation as a single-objective optimization problem.

#### 2) Stationary Points of Potential Game

The Nash equilibria of potential games, referred to as *stationary points* in this paper, are the directionally-local minima of the potential function [13, Prop. 12.2]. The stationary points **W**\*, **K**\* are determined by solving 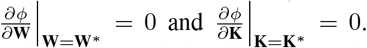 These are points where the brain and the decoder reach steady-state, or are no longer adapting, in the BMI model.

### D. Gradient-Based Co-Adaptation

The brain and the decoder parameters, **W** and **K**, are updated according to the gradient-based adaptation scheme:

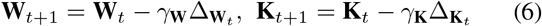

where *γ*_**W**_t__, *γ*_**K**_t__ > 0 denote learning rates and Δ_**W**_t__, Δ_**K**_t__ denote (approximations of) gradients of the agents’ cost functions with respect to their parameters. Note that (6) models adaptation from trial-to-trial; no longer-or shorter-term learning processes are represented. We analyze convergence of (6) to stationary points in two different regimes: 1) *deterministic* gradients; and 2) *stochastic* gradients.

#### 1) Deterministic gradient descent

If learning rates are constant and agents adapt using their exact gradients,

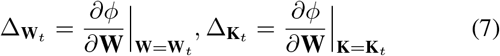

then (6) defines a deterministic dynamical system. To assess convergence, we linearize this system about the stationary points from the preceding section and evaluate eigenvalues of the Jacobian matrix. Since the system is discrete-time, the magnitude of these eigenvalues determines how quickly the co-adaptation dynamics converge to stationarity.

#### 2) Stochastic gradient descent

As noted in [12], evaluating the exact gradients in (7) requires each agent to know the other’s parameters. As an alternative, we consider the dynamics obtained when agents use a *stochastic approximation* of their gradient. One such approximation is [17, Eq. 31].

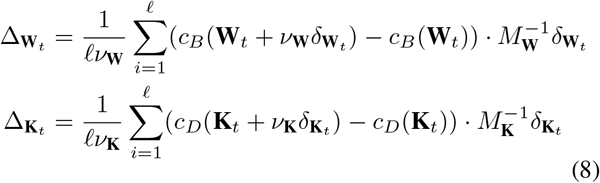

where 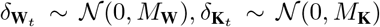 are zero-mean normal random vectors with covariances *M_**W**_, M_**K**_*, respectively. *ν*_**W**_ and *ν*_**K**_ are the difference parameters that scale the contribution of *δ*_**W**_ and *δ*_**K**_, respectively. Note that (8) defines unbiased estimates for the exact gradients in (7) that each agent can evaluate using its own cost and parameters without requiring it know the cost or parameters of the other.

## III. RESULTS

Although the methods described in the preceding section apply to task and neuron spaces of arbitrary dimension, in the present paper we only report results in the special case where the task is 1-dimensional (*d* =1) and there is a single neuron (*n* = 1); generalization of these results to arbitrary dimensions will be the subject of a future study.

### 1) Stationary Points of the Potential Game

When the same value *λ*_**W**_ = *λ*_**K**_ = λ ∈ (0, *τ*^2^) is chosen for the regularization parameters in the brain and decoder cost functions (4), the potential function (5) has two stationary points (the Nash equilibria of the potential game),

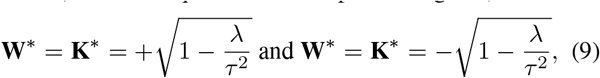

obtained by applying the procedure described in Sec. II-C.2. (When *λ* > *τ*^2^ the only stationary point is at the origin **W**\* = **K**\* =0; when *λ* < 0 there are no stationary points.) Substituting the stationary point (9) into the cost functions (4) yields stationary costs

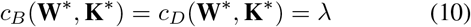

and stationary task error

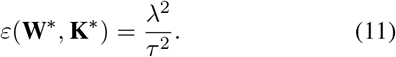

Note that, as the penalty parameter *λ* goes to zero, the stationary task error goes to zero at a quadratic rate.

### 2) Convergence of Deterministic Gradient Descent

When the same value 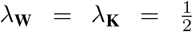 is chosen for the regularization parameters in the brain and decoder cost functions (4) and the task space is normalized so that *τ*^2^ = 1, the eigenvalues of the linearization of (6) at either of the stationary points (9) are

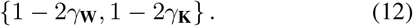

The co-adaptation dynamics converge if the eigenvalues lie inside the unit circle, which occurs precisely when

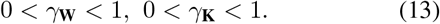

Since the eigenvalue with the maximum magnitude determines the asymptotic convergence rate, the learning rates 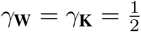 yield the fastest convergence rate.

### 3) Convergence of Stochastic Gradient Descent

To corroborate analytical results reported in the preceding sections, we ran batches of numerical simulations of the co-adaptation dynamics in (6) using the stochastic gradient estimate in (8). We found that these simulations converged to the stationary points in (9) as predicted (Fig. 2).

**Fig. 2.**
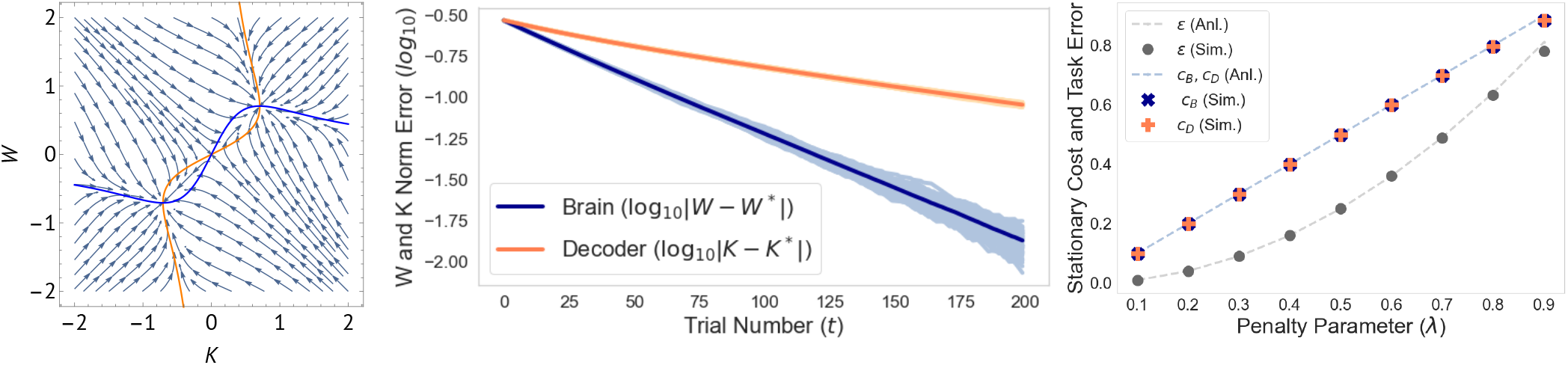
(left) Gradient field of potential function (5) with penalties *λ*_**W**_ = *λ*_**K**_ = 0.5 in (4) and normalized task (*τ*^2^ = 1) as in Sec. II-C.2. Blue and orange curves are *nullclines* (points where 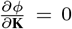 and 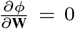, respectively) that intersect at stationary points. (middle) Numerical simulations of co-adaptation (6) with stochastic gradients (8) demonstrate exponential rate of convergence to stationarity (log |**K**_t_ — **K**\* |, log |**W**_t_ — **W**\* | converge to zero at linear rate); the difference in convergence rate is caused by a difference in learning rates (*γ*_**W**_ = 5 × 10^-3^, *γ*_**K**_ = 2.5 × 1θ^-3^). (right) Stationary cost (10) and task error (11) determined analytically (lines) and computed from batches of numerical simulations (markers) for different values of penalty parameter *λ*_**W**_ = *λ*_**K**_ = λ.

## IV. DISCUSSION

### A. Interpretation of Results

#### 1) Stationary points

The formulation of brain-decoder co-adaptation in a game-theoretic framework naturally lead to the consideration of stationary points (Nash equilibria). Our analysis focused on how the penalty parameters *λ*_**W**_, *λ*_**K**_ in (4) affect these stationary points. Focusing on the case where *λ*_**W**_ = *λ*_**K**_ = *λ* ∈ (0, *τ*^2^), we found that *λ* ∈ (0, *τ*^2^) is necessary for the existence of non-trivial stationary points: no points are stationary when *λ* < 0 and only the origin is stationary when *λ* > *τ*^2^. Since these stationary points are our model’s surrogates for the *stable cortical maps* that experiments suggest are highly desirable for skillful control, this finding indicates that penalty parameters play a critical role in ensuring such a stable outcome can be achieved.

#### 2) Convergence of co-adaptation

Assuming both the brain and decoder adapt their parameters using a gradientbased scheme, we analyzed convergence of co-adaptation to stationarity. Our analysis showed that the learning rates *γ*_**W**_,*γ*_**K**_ play a critical role in convergence. Focusing on the case where 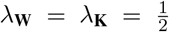 and *τ*^2^ = 1, we found that convergence is guaranteed so long as

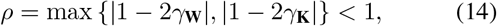

in which case *ρ* determines the rate of convergence since **W**_*t*_ → **W**\* and **K**_t_ → **K**\* exponentially in time/trial *t* with rate *ρ* < 1:

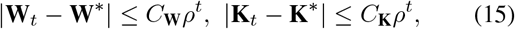

for some fixed constants *C*_**W**_, *C*_**K**_ > 0. Our findings demonstrate that the decoder’s learning rate must be chosen carefully, as learning too fast may lead to an unstable outcome, whereas learning too slow may limit the rate of convergence to stationarity. These analytical results are corroborated by previous simulation and experimental findings [6], [12].

### B. Future Directions

In this paper, we have detailed an initial theoretical framework that can be used to inform adaptive decoder design for shaping brain learning and optimizing BMI performance. The dynamic game formulation of a co-adaptive BMI can be extended to BMI models and experiments where the decoder and brain may have separate objectives, which may be necessary for the decoder to guide the brain’s learning. Our stochastic simulations confirmed the deterministic analysis and will allow us to evaluate co-adaptive BMI scenarios that are less mathematically tractable. Example of such mathematically difficult scenarios include: (a) non-stationary costs, (b) learning rates that are dependent on time or task, (c) nonlinear thresholds for firing rates. Next steps to guide the BMI model presented here will be a comparison to experimental data of co-adaptation [7].

